# The Cognitive Mechanisms That Drive Social Belief Updates During Adolescence

**DOI:** 10.1101/2020.05.19.105114

**Authors:** I. Ma, B. Westhoff, A.C.K. van Duijvenvoorde

## Abstract

Adolescence is a key life phase for developing well-adjusted social behaviour. Belief updates about the trustworthiness of peers are essential during adolescence as social reorientation emerges and peer relationships intensify. This study maps the age-related changes of those belief updates during adolescence (*n* = 157, 10-24 years). We used computational modelling and an information sampling paradigm to reveal that three cognitive mechanisms contribute to age-related changes in those belief updates: prior beliefs, prior uncertainty, and uncertainty tolerance. The age-related changes in these three cognitive mechanisms result in increasingly adaptive belief updates from early to mid-adolescence when it comes to beliefs about trustworthiness. Our findings shed light on age-related changes in adaptive learning about others during adolescence.

---

Adolescence is a life-phase accompanied by a strong social reorientation in which achieving and upholding a positive peer status becomes a salient social goal^1–5^. To reach this social goal, adolescents’ beliefs about the trustworthiness of a peer is especially important, as trusting the right person is essential to build positive social interactions, such as friendships and romantic relationships^6,7^. However, trusting the wrong person may have harmful consequences (e.g., social rejection, gossiping about entrusted secrets) to which adolescents are highly sensitive^8,9^. Adjusting our beliefs about others based on outcomes of social interactions is critical for successful navigation of the social world. We hypothesize that three distinct cognitive mechanisms show age-related change during adolescence and give rise to age-related changes in their belief updates: 1. the prior beliefs (our expectations of others trustworthiness), 2. the uncertainty about those prior beliefs, and 3. how much uncertainty the individual finds tolerable (uncertainty tolerance). These mechanisms are not directly observable and therefore challenging to expose with empirical measures alone. We use Bayesian computational models combined with an information sampling paradigm to distinguish between these mechanisms and uncover their contribution to age-related changes in belief updates during adolescence.

We first consider age-related changes in prior beliefs. Applied to trust, prior beliefs are the expectations that adolescents have about the trustworthiness of (unknown) others. Past empirical studies show that adolescents become more trustworthy with age and trust others more, which stabilizes into adulthood^10,11^. This suggests that during adolescence, priors may shift to believing that others are more trustworthy compared to beliefs during childhood. An individual’s prior belief influences their belief updates. That is, new information can confirm a prior belief, making the updated belief (posterior belief) stronger (i.e., more certain), or disconfirm the prior belief then the posterior shifts (e.g., “peers are more trustworthy than I first thought”).

The uncertainty of prior beliefs is the second cognitive mechanism in which we consider age-related changes. Consider the following example to illustrate prior uncertainty: someone with low prior uncertainty could believe that all peers are mostly trustworthy and expect little variation between individual trustees. A person with high prior uncertainty might expect the same level of peer trustworthiness on average but also expects a large difference between individual trustees. Evidently, strong priors are not very useful after the environment has changed. Bayesian models indeed show that prior uncertainty is modulated by the volatility of the environment and affect the strength of belief updates^12^. A change in the environment increases prior uncertainty, which leads to heightened sensitivity to new information^13^. Uncertainty about the environment is theorized to contribute to the onset and duration of developmental sensitive periods of learning^14,15^. We apply this reasoning to adolescence. Specifically, adolescence changes the set of displayed social behaviours and increases their frequency (e.g., courtship or competitive behaviour such as gossiping). In addition, transitioning from primary to high school exposes the adolescent to new peer groups and increases uncertainty about the generalizability of previously learned priors over social behaviours (e.g., “childish” games such as playing tag may not be socially accepted anymore in high school). We therefore hypothesize an *increase* in the uncertainty of prior beliefs about the trustworthiness of others during adolescence.

Third, we consider changes in uncertainty tolerance during adolescence. Trusting is a decision under uncertainty, as the exact likelihood of reciprocation is unknown (this uncertainty is sometimes called ambiguity in economics). Past studies using lottery tasks with unknown outcome probabilities suggest that uncertainty tolerance decreases from adolescence to adulthood^16,17^. One study found that adolescents are more uncertainty tolerant than children and adults^18^ (but see^19,20^). It is unknown if the age-related changes in uncertainty tolerance found in lottery tasks generalizes to social contexts such as trust. We explore whether age-related changes in uncertainty tolerance drive adolescents’ belief updates about others’ trustworthiness.

Taken together, here we examine the age-related changes in prior beliefs, prior belief uncertainty, and uncertainty tolerance as underlying mechanisms in trustworthiness belief updates. These mechanisms are internal and not directly observable but can be exposed using active sequential information sampling and computational modelling. To this end, we used the Information Sampling Trust Game and computational models^21^ in an adolescent sample with a wide age range (10-24 years). These findings shed light on the cognitive mechanisms that drive social belief updates during adolescence.

## METHODS

### Participants and experiment procedure

A total of 157 adolescents (of which 75 boys) completed the experiment (range = 10-24 years, *M* = 17.50, *SD* = 4.34). These were uniformly sampled across 5 age bins. Participants were screened for colour blindness, psychiatric and neurological disorders, IQ was estimated by using subtests of the WISC and WAIS. IQ scores fell in the normal range (*M* =107.5, *SD* = 10.9, range = 80-135), and did not correlate with age (*r*_*s*_ = 0.119, *P* = 0.138), parental social economic status (SES) was estimated by highest educational attainment of the caregiver(s). SES also did not show a relationship with age (Kruskal-Wallis rank sum returned *χ*^2^(4, *n* = 157) = 6.342, *p* = 0.175). The institutional review board of the Leiden University Medical Center approved this study. Written informed consent was given by adult participants, and by parents in the case of minors (minors provided written assent). This behavioural study was part of a larger imaging study, not relevant to the current study. All participants performed the Information Sampling task in a quiet room near the neuroimaging labs of Leiden University. The task took approximately 30 minutes to complete.

### Information Sampling Trust Game Task

The Information Sampling Trust Game (Figure 1A) combines a well-established behavioural economics paradigm known as the single-shot Trust Game^7^ with sequential information sampling. On each trial there are 2 players: the investor and a trustee. All our participants played the task in the investor role. At the beginning of each trial, participants were endowed with 6 tokens which they could invest (entrust) in the trustee. Participants were told trustees previously participated in a different experiment where they made decisions to either reciprocated or keep entrusted money. If invested, the trustee received the endowment multiplied by 4 (24 tokens) and subsequently decided to either reciprocate, resulting in 12 tokens for both players, or defect by keeping all 24 tokens leaving 0 tokens for the participant. Participants were also given the option of not investing by keeping the initial endowment (6 tokens) (see Figure 1A, payoff matrix). Participants were told that the trustees previously completed 25 trials (decisions) in the trustee role played with different investors and that their decisions to either reciprocate or not were stored in a covered 5×5 grid. On each trial, participants were given the opportunity to sequentially sample information about the trustee’s reciprocation history before deciding to either invest or not invest. They sampled by sequentially turning tiles in the grid (turn minimum 0 and maximum 25 tiles per trial). If the tile turned green, the trustee had decided to reciprocate the trust of the past investor, while red indicated a past decision to defect. It was also clarified that the location of the tile was not informative, that each trial would be played with a new unknown trustee, and the ratio green to red tiles would therefore vary a lot between trials. Unbeknownst to participants, grid outcomes were computer-generated and drawn from the following probabilities: 0.0, 0.2, 0.4, 0.6, 0.8, and 1.0, where 0.0 is completely untrustworthy (all red) and 1.0 is fully trustworthy (all green). Each subject sampled information about 60 different trustees (trials), resulting in a maximum of 1500 sampling decisions. There were no explicit sampling costs other than the time and effort involved in turning tiles. Participants were fully informed of the payoff matrix. The outcomes of trust decisions were not shown during the task to avoid meta-beliefs about the reliability of the acquired information. Instead, participants were told that 3 trials would be randomly selected at the end of the task and their average amount of tokens would be converted to money and paid to the participant (Supplement).

**Figure 1.**
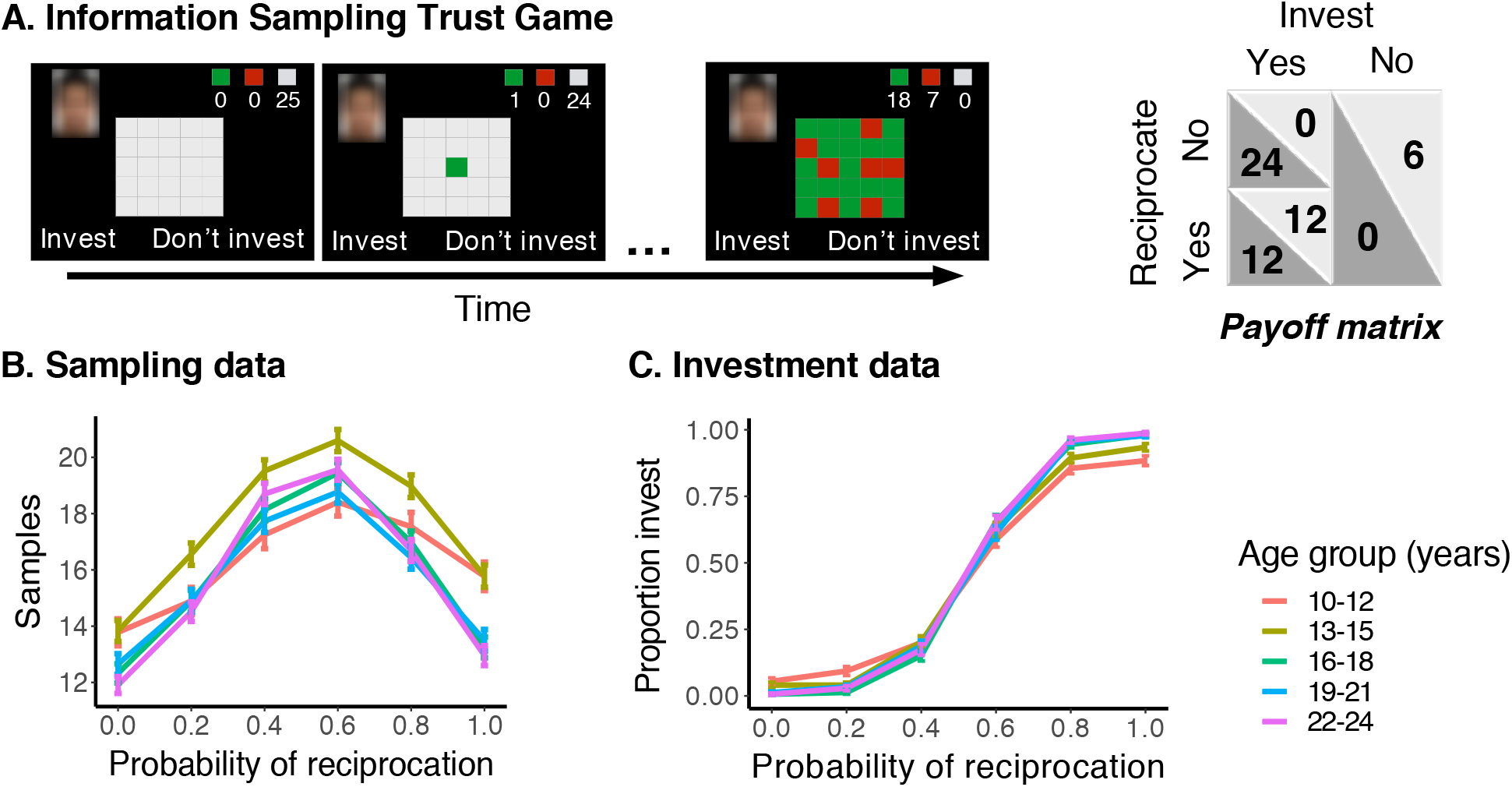
Information Sampling Trust Game and data. **A.** Task trial sequence example and payoff matrix. The participant could sample a trustee’s reciprocation history with other investors up to 25 times by turning tiles in a 5 by 5 grid. Green = reciprocated trust, red = betrayed trust, grey = not sampled. Investment outcomes were not shown during the task. Six reciprocation probabilities (*r* = 0.0, 0.2, 0.4, 0.6, 0.8, 1.0) generated the outcomes in the grid. **B.** The number of samples (mean and standard error of the mean (s.e.m.)) as a function of reciprocation probability per age group (years). This shows that participants sampled more when the outcomes were least conclusive, i.e. when the reciprocation probability was closest to 0.5. Early-adolescents (10-12 years) showed less variation as a function of reciprocation probability compared with other ages (i.e., a more flattened sampling distribution across reciprocation probabilities). **C.** Proportion of investments as function of the generative reciprocation probability for each age group. This shows that investment decisions follow the generative reciprocation probability but that this effect is more pronounced in older adolescents. With age, adolescents trusted more often when the trustee was highly trustworthy (reciprocation probability = 0.8 and 1.0) and less when the trustee was untrustworthy (reciprocation probability = 0.0).

### Computational model

*The Uncertainty model* is based on the concept of sampling to reduce uncertainty until a subjective uncertainty tolerance criterion is met. The model consists of four components: a prior belief over the reciprocation probability (*r*), an evolving posterior distribution over *r*, an uncertainty tolerance criterion, and decision noise. As explained in the introduction, individuals with different prior belief distributions will also have different posterior distributions after they observe the same sample outcomes (Figure 2).

**Figure 2.**
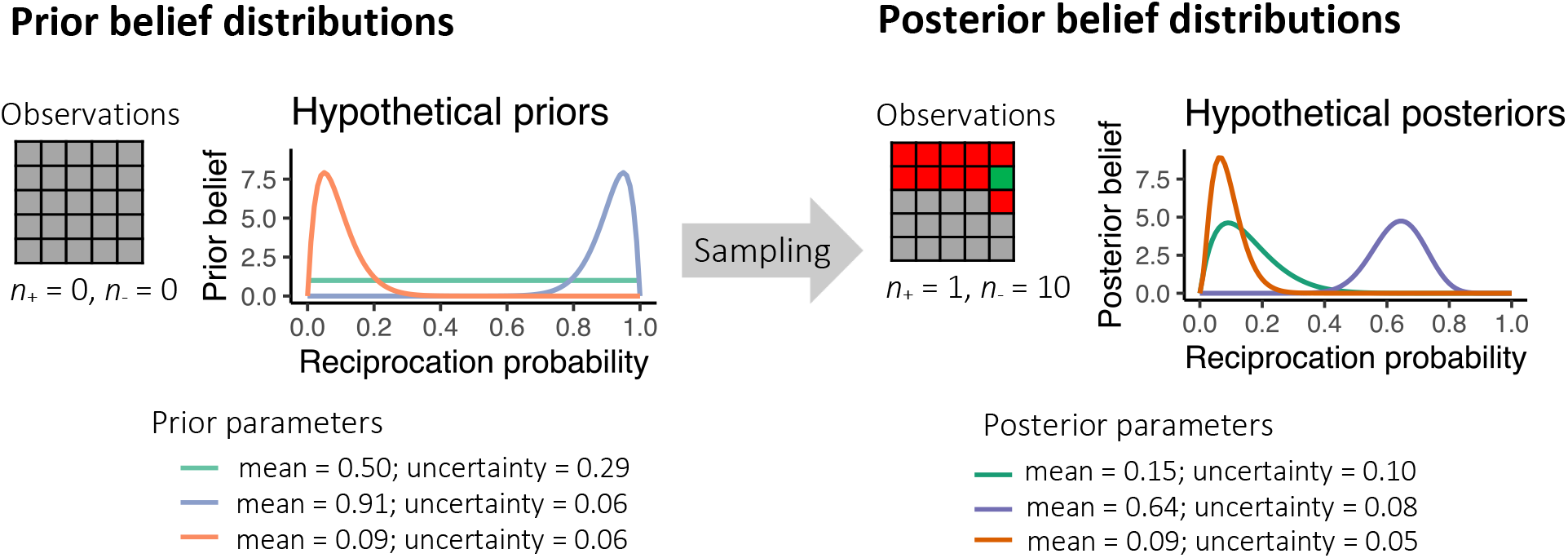
Illustration of how prior belief distributions update after observing 10 red tiles (defect) and 1 green (reciprocation) tile. The orange, green and blue lines represent three hypothetical subjects’ prior distributions and their corresponding posterior distribution. These three hypothetical subjects were selected to show that the posterior beliefs can be quite different depending on the mean and uncertainty of the prior beliefs. The orange prior distribution reflects a belief that lower reciprocation probabilities are more likely. The observed outcomes exactly match that prior belief. The posterior uncertainty therefore decreases but the posterior mean does not update. The green prior distribution has a maximal uncertainty, i.e., a belief that all reciprocation probabilities are equally likely. The posterior then shows a both large update in the mean and a reduced uncertainty. The blue prior distribution reflects a belief that higher reciprocation probabilities are more likely. The sample outcomes disconfirm this belief. The posterior then shows a large update and becomes more uncertain.

Formally, in our model the *state* is defined by the number of turned green tiles (*n*_+_), and the number of turned red tiles (*n*_-_), and the *actions* are to either sample or stop sampling until all 25 tiles are sampled. Specifically, the model assumes that people do not know the trustee’s exact trustworthiness by using a Bayesian belief distribution over the possible range of *r*. The conjugate prior belief distribution over *r* is a beta distribution with parameters *α*_0_ and *β*_0_. As information is sampled, the evolving posterior distribution is:

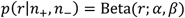

where *α* = *α*_0_ + *n*_+_ and *β* = *β*_0_ + *n*_-_.

As shown in Figure 2, the more uncertain a prior belief is, the more a new sample will reduce that uncertainty. In addition, if a sample is highly consistent with the prior belief, the posterior mean will shift less than when a sample disagrees with the belief^12^. Uncertainty of the belief is operationalized using the standard deviation of the posterior distribution:

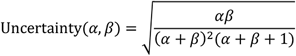

As sampling decreases uncertainty and at some point, the uncertainty will reach the subject’s uncertainty tolerance criterion *k.* This criterion reflects how much uncertainty is tolerated by the subject. As the uncertainty reduces and approaches this criterion, the probability that the subject takes another sample becomes smaller. These probabilities are given through the softmax function, allowing for decision noise *τ*:

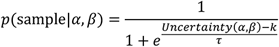

Where a larger *k* reflects more uncertainty tolerance and a larger *τ* reflects more decision noise.

We fitted the model using a log likelihood optimization algorithm as implemented in the fmincon routine in MATLAB (Mathworks) using 100 combinations of starting points to avoid local minima. Four free parameters are fitted for each subject: *α*_0_, *β*_0_, *k,* and *τ*. We used between-subjects Bayesian Model Selection^22,23^ to assess variation in the best fitting model between different ages using five age groups.

### Alternative models

We also considered two families of alternative models that did not fit as well as the Uncertainty model (formal descriptions in Supplement): The *Sample Cost model*, which uses the Bayesian belief distribution to compute the normative solution for every state. The *Threshold model* is a heuristic model that does not use Bayesian beliefs distributions. For model comparisons we calculated the difference between model evidence in terms of BIC for each model pair for each subject. To assess the significance, we used bootstrapping to compute the 95% confidence intervals of the summed difference in BIC using 10^5^ iterations.

## RESULTS

### Descriptive statistics

On average participants sampled 16.229 (*SD* = 7.532) of 25 times per trial. We used a mixed-effects model (LME4 in R^24,25^) to test whether the consistency of sample outcomes (information consistency) indeed affected the probability of sampling, as was expected based on non-social sampling studies^26^ (Supplement). The results showed that participants were more likely to draw another sample when the information consistency was lower (i.e., smaller difference between green and red outcomes) (β = 0.830, *P* < 0.001). This effect interacted with the linear effect of age (β = 0.174, *P* < 0.001), and the quadratic effect of age (β = −0.036, *P* = 0.002; see Table S2.1 for full model results). This shows that participants were generally more likely to turn over another tile when outcomes were inconclusive. This effect became stronger with age, especially during early-adolescence (i.e., 10-12-year-olds) (Figure 1B).

Next, we used a mixed-effects model to examine whether the invest decisions (i.e., decisions to trust or not) were predicted by the reciprocation probability and whether this differed across age (Supplement). Participants invested more frequently with higher reciprocation probabilities (Figure 1C), although this effect interacted with age (age linear: β = 2.123, *P* < 0.001; age-quadratic β = −0.906, *P* < 0.001). Post-hoc analyses per reciprocation probability showed that for high reciprocation probabilities, the number of decisions to trust increased from early to mid-adolescence (Supplement Table S2.2). This also led older adolescents to have a somewhat higher average expected reward compared to younger adolescents. There were no significant age-related changes in trust decisions for reciprocation probabilities closer to 0.5 (all *P* > 0.071) (Figure 1C; Table S2.3).

### Computational modelling results

The Uncertainty model fitted the data better than the two alternative models: The Threshold and the Sample Cost models (Figure 2). Bootstrapping the 95% CI’s showed that this difference was significant (Figure 2B). Moreover, Spearman correlations showed no significant relation between the BIC difference and age (Supplement Table S1.2). Importantly, this suggests that out of these three models, the Uncertainty model was the best fitting model for all ages (see Figure 2C for model fits per age group, also see Supplement Table S1.2).

**Figure 2.**
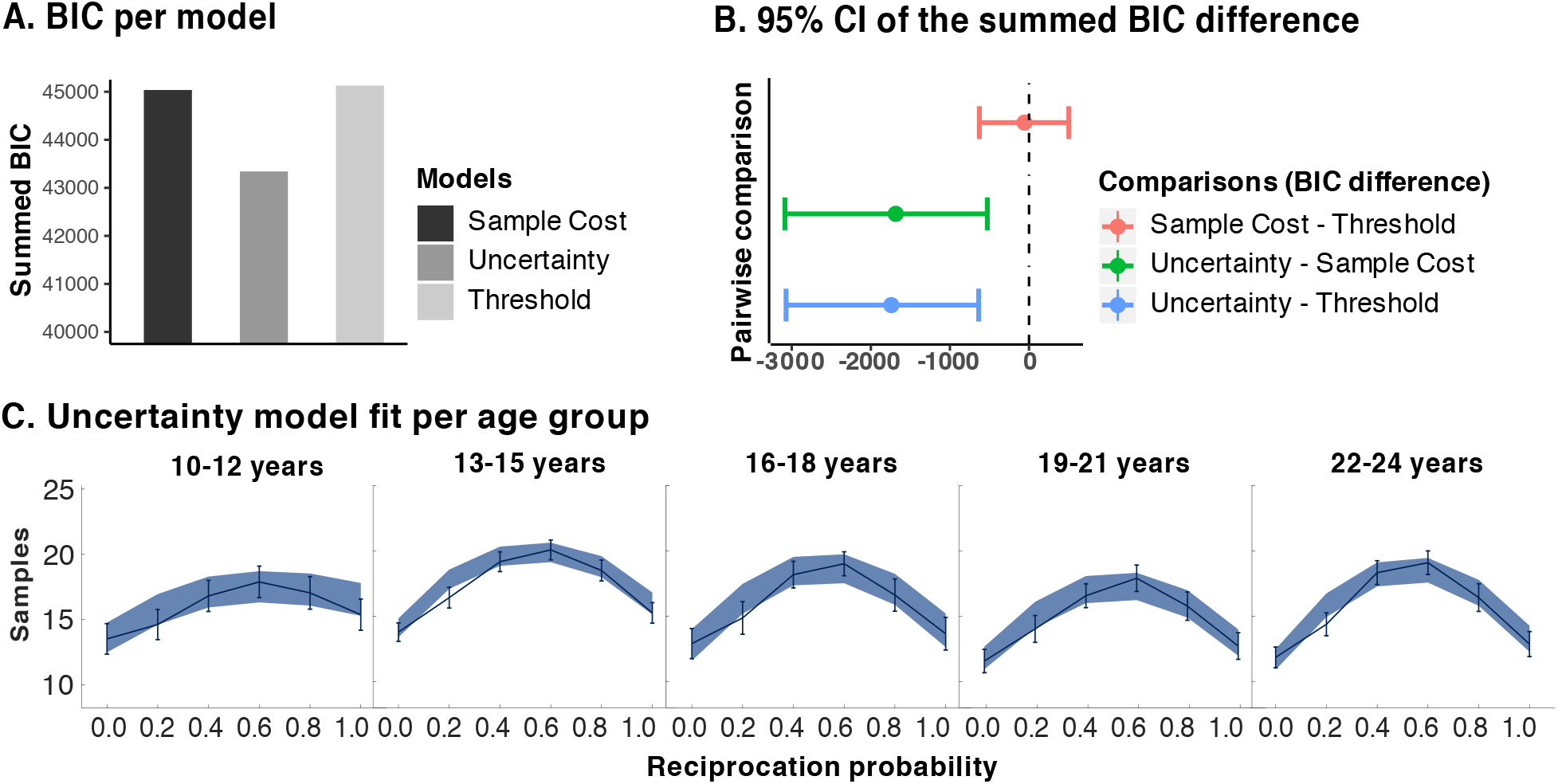
Model fit results. **A.** The summed Bayesian Information Criterion (BIC) score per model. Lower BIC values indicate a better fit, thus showing that the Uncertainty model fits best. BIC scores were computed for each participant and each model. **B**. 95% confidence intervals (CI) of the summed BIC difference between models. Zero indicates no difference between models. Negative values are in favour of the model before the subtraction sign, as lower BIC indicates a better fit. The Uncertainty model fits significantly better than the Sample Cost and Threshold models (95% CI does not contain zero). The BIC scores of a model pair were subtracted from each other for each participant, thereby obtaining one difference score per participant for each model pair. To assess significance, the 95% confidence interval of the BIC difference was computed using bootstrapping with 10^5^ iterations. **C.** Winning model fit across age. The line graph represents the mean and s.e.m. of the raw data. The shaded area is the s.e.m. of the model fit. This shows that the Uncertainty model fitted well for each age group.

### Age-related changes in Uncertainty model parameter estimates

Adding quadratic age term to the linear regression model only improved the fit for the relation between age and prior uncertainty parameter estimates. For all other parameter estimates the regression model fits did not improve and we therefore report only linear age effects for those parameters (see Supplement for the model improvement tests and Bonferroni-Holm correction). The *mean of the prior beliefs* showed a significant change with age (prior distribution mean = 0.450, *SD* = 0.154; age linear, β = 0.212, *P* = 0.009. With age, participants expected trustees to be more trustworthy, though this age effect was subtle (Figure 3). The *uncertainty of the prior beliefs* showed that prior beliefs became more uncertain during adolescence, the quadratic effect suggests that this age-related change was stronger from early to mid-adolescence (ages 10-16 years, Figure 3; age linear, β = 0.027, *P* < 0.001; age quadratic, β = 0.020, *P* = 0.005). The *uncertainty tolerance criterion* estimates increased linearly with age (β = 0.012, *P* < 0.001, Figure 3), suggestion that adolescents monotonically became more uncertainty tolerant with age. Finally, we found that *decision noise* (inverse temperature estimate) did not change with age (β = 0.11, *P* = 0.600). This shows that the degree to which participants followed the fitted Uncertainty model predictions did not change with age. More specifically, the degree to which the difference between the uncertainty and the uncertainty tolerance criterion mapped onto participants’ sampling probabilities did not change with age. This latter finding is another indicator that the model fit did not significantly differ with age and therefore gives more confidence in the interpretability of the estimates of our parameters of interest as function of age.

**Figure 3.**
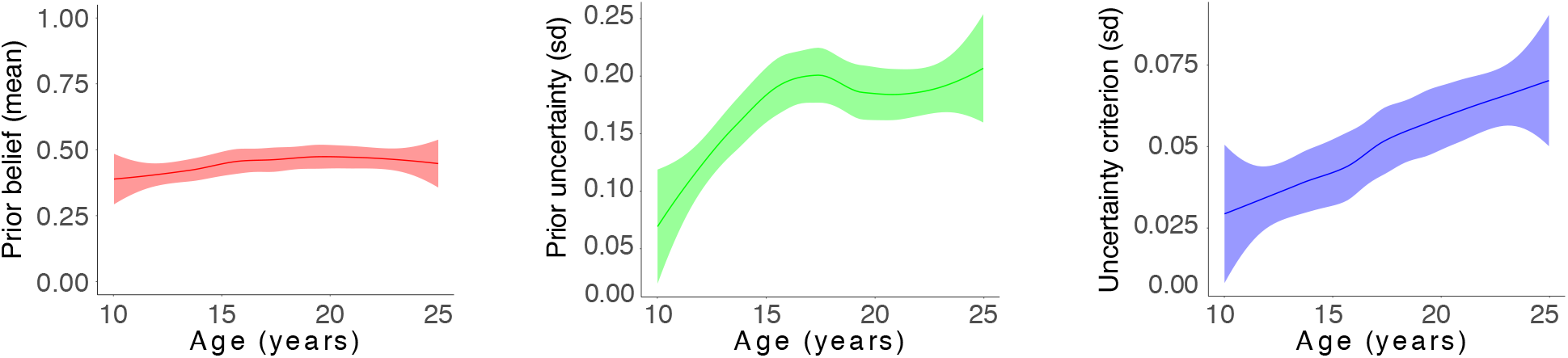
The winning Uncertainty model parameter estimates. Uncertainty model prior mean, prior uncertainty, and the uncertainty criterion as a function of age (using Loess method). Note that a uniform prior corresponds to a standard deviation of 0.289, thereby creating an upper bound for the prior uncertainty. Lower prior uncertainty values reflect a more certain prior.

In summary, we show that the prior beliefs, uncertainty about these prior beliefs, and uncertainty tolerance changed from early to mid-adolescence. While the prior belief change was subtle, the uncertainty in these prior beliefs more strongly and rapidly increased from early to mid-adolescence. In addition, uncertainty tolerance increased more slowly and monotonically with age. We verified through model recovery that the models were distinguishable and through parameter recovery that the number of trials was sufficient to accurately estimate parameters (Supplement).

### No difference in prior parameters in first and second task-half

To test whether sampling information about trustees during the task had changed participant’s prior parameter estimates, we refitted our model parameters to the first half and second half of the task separately. We then assessed the stability of prior parameter estimates by comparing the prior estimates *α*_0_ and *β*_0_ between the task halves. The Wilcoxon sign-rank tests showed difference between task halves in the prior parameter estimates (*α*_0_ prior estimate, *z* = 1.406, *P* = 0.160, median difference < 0.001, 95% CI [-0.122, 0.315]; *β*_0_ prior estimate, *z* = 1.133, *P* = 0.257, median difference < 0.001, 95% CI [-0.478, 0.465]). Moreover, Spearman-rank correlations showed no significant relation between age and the difference in prior parameter estimates between the first and second task halves (*α*_0_ prior estimate *r*_*s*_ = −0.046, *P* = 0.565; *β*_0_ prior estimate *r*_*s*_ = −0.058, *P* = 0.573). This suggests that the findings in our study regarding the prior estimates were not likely confounded by participants learning the trustworthiness distribution during the task or age-related changes therein.

## DISCUSSION

We identified cognitive mechanisms that underlie adolescents’ updates in belief about the trustworthiness of others and mapped the age-related changes in these mechanisms from early to mid-adolescence. As hypothesized, the three cognitive mechanisms that showed age-related changes were prior beliefs, the uncertainty of prior beliefs, and uncertainty tolerance.

Perhaps the most striking finding were the age-related changes in the uncertainty of the prior beliefs (i.e., the uncertainty about expectations about others behaviour). Early-adolescence was mainly characterized by a rapid increase in the uncertainty about others trustworthy behaviour. This increase in uncertainty introduces that adolescents gradually relied less on their prior beliefs and relied more on their sampled evidence. The increase in prior uncertainty may be adaptive by allowing mid-adolescents to more efficiently learn that some peers are highly trustworthy while others are not. More generally speaking, these findings frame adolescence as a developmental phase during which people start to consider a broader hypothesis space about the trustworthiness of others. This idea is corroborated by recent empirical findings showing that adolescents are more flexible learners than adults in the social domain^27^, and by theoretical work suggesting that changes in the (social) environment result in a sensitive period for learning and exploration^14,15^. We also found that people expected others to be slightly more trustworthy this led, early-adolescents to sample somewhat more when the reciprocation probability was above 0.5 compared to when it was lower and trusted fully trustworthy trustees less often than older adolescents. In light of these previous empirical and theoretical studies, our findings set the stage for early to mid-adolescence as a crucial developmental phase of social learning through adjustments of uncertainty in prior beliefs.

In addition, we found that adolescents became more uncertainty tolerant with age. This is contrary to our expectations, and to previous findings in non-social economic decision tasks, in which ambiguity about the probability of outcomes is manipulated. In most of these studies, ambiguity tolerance decreases (or even peaks) in adolescence compared to adulthood. The current findings seem to indicate that uncertainty tolerance in a social context may follow a distinct developmental trajectory, which should be confirmed in future studies. Nonetheless, uncertainty tolerance in social contexts should be interpreted in combination with the prior parameters to explain the participants data. That is, in our study a low uncertainty tolerance, combined with a low prior uncertainty about others as seen in 10-12-year-olds does not allow these early-adolescents to adapt as well to different levels of trustworthiness of others. On the other hand, 13-15-year-olds were increasingly uncertainty tolerant, but also more uncertain about other’s trustworthy behaviour. As shown in Figure 1B, this resulted in more sampling overall (i.e., more information seeking regarding others behaviour) and a stronger adaptation to the specific reciprocation probabilities compared to younger ages. Taken together, our findings demonstrate that age-related changes in belief updates can be explained by the three underlying cognitive mechanisms that are captured by our model.

We found no significant age-related changes in decision noise. In our study, decision noise reflects the degree to which sampling decisions are driven by the difference between uncertainty and the uncertainty tolerance criterion (i.e., high decision noise means that people make their decisions to either sample or stop more randomly). This is consistent with previous studies which did not find age-related changes in random decision-making^28,29^. Instead, previous studies suggest that younger children show higher decision noise if it serves exploratory purposes (for review see^30^). It should be noted that decision noise in our study does not reflect exploratory behaviour because stopping in this paradigm does not result in exploration. The fact that we did not find age-related differences in decision noise in the sampling decisions, further validates the assessment of age-related changes in the model parameter estimates.

Moreover, the Uncertainty model fitted well and performed best for all ages. We specifically ruled-out a family of normative models (i.e., Sample Cost model variants) and heuristic models that were not based on Bayesian belief distributions (Threshold model variants). In many reinforcement learning and decision-making tasks, winning models differ between age groups, which obstructs generalization of model comparisons in studies on adults to other age-groups^31^. For example, adolescents show different reinforcement learning strategies compared to adults by not benefitting from counterfactual feedback^32^. In a different context, children more often use model-free strategies in a reinforcement learning task whereas model-based choices became more frequent in adolescence and adulthood^33^. Our results show that people use the same strategy to sample information about others trustworthiness across adolescence, but that specific parameter estimates within that model are subject to age-related changes. Computational modelling therefore allowed us to dissociate between cognitive strategies and uncover age-related changes in cognitive processes that could otherwise not be revealed.

Exploring the effects of peer status on trustworthiness information sampling was beyond the scope of this study but is a potentially promising avenue for future research. Responses to adverse social events such as social exclusion depend on relative peer status^34^. For example, children and adolescents who are chronically rejected by peers show heightened neural sensitivity to acute events of exclusion by peers^35,36^. These rejected children and adolescents may have different priors over the trustworthiness of others than their accepted peers (e.g., those with experience of frequent rejection may have a prior belief that others are untrustworthy). A potentially interesting consequence of such individual differences in prior beliefs about trustworthiness is that it can further reinforce individual differences in beliefs about the social environment due to stochastic outcomes. Specifically, in real-life, reciprocity is not always in full control of the trustee: sometimes unintended situations can cause hinderance of trust reciprocations (e.g., accidentally dropping someone’s phone causing it to break). These negative outcomes can even occur when the trustee is actually trustworthy. Someone with a strong prior belief about others being untrustworthy would likely stop sampling sooner and not trust when they encounter these negative outcomes than someone with neutral or positive priors. By stopping to sample (i.e., avoidance), the strong belief about people being untrustworthy will never be updated to the true value (e.g., people may be more trustworthy). This avoidance mechanism based on initial impressions can therefore lead to biased beliefs about the statistics of the environment^37^. It is important to understand individual differences in prior beliefs about others and explore potential ways to counteract those, for example, by reducing the opportunities to avoid^38^. It is especially relevant to understand how these mechanisms develop across adolescence, as our findings indicate that this is a developmental phase during which beliefs about the social environment are adjusted.

The potential applications of our approach further extend to developmental psychiatric disorders. For example, obsessive compulsive disorder and high levels of compulsivity are associated with aberrant non-social information sampling cost functions^39,40^. In addition, previous studies used behavioural economic games such as trust games to reveal aberrant social decision-making psychiatric disorders^41–43^, including autism spectrum disorder^44^, borderline personality disorder^43,45,46^, and ADHD^47,48^. However, how people actively sample and use information to initiate social interactions and how this may depend on prior beliefs is an underexplored topic in these fields. Our study provides insights into the underlying process of these decision-making strategies in typical adolescents and offers a benchmark to understand atypical development.

