## Supplement for "The Cognitive Mechanisms That Drive Social Belief Updates During Adolescence"

### ONLINE SUPPLEMENTARY MATERIALS

#### Participant bonus fee

Unbeknownst to the participants, payoff was not dependent on trustee decisions, instead three trials were selected and their outcome was averaged to determine payoff. If invested and the generative reciprocation probability was > 0.5 then the outcome was 12 tokens, and 0 tokens if < 0.5. To avoid extreme bonus differences between participants, the payoff trial selection procedure was not fully randomized such that each participant ended with a task performance bonus about between 3 and 9 tokens. The tokens were converted to money as follows: 3 = €1, 4 = €2, 5 = €2, 6 = €3, 7 = €3, 8 = €4, and 9 = €5 and this amount was added to the participation fee.

#### Computational models

Note that for all models, we tested different variants within each model, and selected the best fitting version for between model comparisons. Here, we denote the best fitting version of each model, for all versions see Ma et al., (2018).

##### *The Sample Cost model*

*The Sample Cost model* uses the Bayesian belief distribution over trustworthiness to compute the expected utility of sampling and stopping for every possible state in the task through forward reasoning. It consists of four components: prior beliefs over the trustworthiness ($r$), an evolving posterior distribution over $r$ iterative maximization of future expected utility under this posterior distribution, and decision noise. The conjugate prior over *r* is a beta distribution with priors $\alpha_0$ and $\beta_0$, and parameters $\alpha = n_+ + \alpha_0$ and $\beta = n_- + \beta_0$, where $n_+$ is the number of green samples and $n_-$ is the number of red samples. The posterior over *r* is:

$$p(r|n_+, n_-) = Beta(r; \alpha, \beta)$$

Investing results in either reciprocation (outcome = 1 with probability *r*, then the investment amount is multiplied by m = 2) or betrayal (outcome = 0 with probability 1 - *r*, then the investment amount is multiplied by m = 0). The agent does not know *r* and therefore has to marginalize over *r*, using the current posterior. This gives the conditional distribution of outcome given $\alpha$ and $\beta$:

$$p(\text{outcome} = 1|\alpha, \beta) = \int p(\text{outcome} = 1|r)p(r|\alpha, \beta)dr = \int rp(r|\alpha, \beta)dr = \frac{\alpha}{\alpha+\beta} \quad (1)$$

The expected utility of *not* investing is $U_0 = 1$. The expected utility of investing, $U_1$, with a free parameter $\lambda$ for risk attitude becomes:

$$U_1(\alpha, \beta) = \mathrm{E}[\text{outcome}|\alpha, \beta] - \lambda \mathrm{Var}[\text{outcome}|\alpha, \beta]$$
$$= \frac{m\alpha}{\alpha + \beta} - \frac{\lambda m^2 \alpha\beta}{(\alpha + \beta)^2} \quad (2)$$

The agent can decide between sampling (a = 1) or stopping (a = 0) at any time except when t = T+1, then all boxes are opened and only stopping (a = 0) is possible. The value of a state-action pair is given by the Bellman equations (Bellman, 1952). Specifically, the expected value of the state $(\alpha, \beta)$ is the higher of the expected utilities of not investing and investing:

$$Q_t(\alpha, \beta; a = 0) = max\{U_0, U_1(\alpha, \beta)\} \quad (3)$$

When t = T+1, the value of the state $(\alpha + \beta)$ is $V_t(\alpha, \beta) = Q_{T+1}(\alpha, \beta;$ a = 0). At earlier times, $V_t$ is the larger of the expected utilities:

$$V_t(\alpha, \beta) = max\{Q_t(\alpha, \beta; a = 0), Q_t(\alpha, \beta; a = 1)\} \tag{4}$$

The expected value of a sampling action (a = 1) at time *t* in the state $\alpha, \beta$ subtracting the subjective cost of a sample *c* is:

$$Q_t(\alpha, \beta;\ a = 1) = \frac{\alpha V_{t+1}(\alpha + 1, \beta) + \beta V_{t+1}(\alpha, \beta + 1)}{\alpha\ +\ \beta} - c \tag{5}$$

By starting with the final state (when *n* = T), we can obtain the Sample Cost solution for every possible state (dynamic programming). In the Sample Cost model, the decision variable (DV), is the difference between the utilities of sampling and stopping:

$$DV(\alpha, \beta) = Q_t(\alpha, \beta; a = 1) - Q_t(\alpha, \beta; a = 0) \tag{6}$$

The Sample Cost policy would be to sample when the DV is positive; however, we introduce decision noise through a softmax function:

$$p(\text{sample}|\alpha, \beta) = \frac{1}{1 + e^{-\frac{DV(\alpha,\beta)-k}{\tau}}}$$

This model version has free parameters: $\alpha_0, \beta_0, c, \lambda, \tau,$ and $k$ which are fitted for each subject.

*The Threshold model*

In the *Threshold Model,* the agent keeps track of the absolute difference between positive and negative information and stops sampling when this difference reaches a bound. However, to be consistent with our other models and with most of the value-based decision literature, we use soft rather than hard bounds. The bound *b* takes three possible values, depending on the sign of $n_+ - n_-$:

$$b = \begin{cases} b_+ & \text{if } n_+ > n_- \\ \frac{b_+ + b_-}{2} & \text{if } n_+ = n_- \\ b_- & \text{if } n_+ < n_- \end{cases} \tag{7}$$

The probability that the agent stops sampling is a logistic function of the difference between this decision variable and a bound *b*:

$$p(\text{sample}|n_+, n_-) = \frac{1}{1 + e^{-\frac{DV(n_+,n_-)-b}{\tau}}} \tag{8}$$

This model version has free parameters: $b_+$, $b_-$, and $\tau$ which are fitted for each subject.

**Within-model comparisons results**

For each model, we first fitted the basic version - the version with the fewest free parameters - and then tested whether adding a free parameter significantly improved the model fit (also see (Ma et al., 2018) for details). The assessment of these additional free parameters was based on the trust game literature. Here, we describe the additionally tested free parameters for each model across all participants.

*Uncertainty model: prior beliefs improve the model fit.* For the same reasons as described earlier, we tested the improvement in model fit when adding a prior belief, also estimated for each subject.

In the basic version of our model, we used an uninformative, uniform prior (i.e., $\alpha_0$ = 1 and $\beta_0$ = 1). In the model version with prior beliefs, we fitted $\alpha_0$ and $\beta_0$ as free parameters for every subject. We found that allowing for a subjective prior improved the model fit (Table S1). The median of individual prior means was $r$ = 0.47 (bootstrapped 95% CI [0.45, 0. 50]).

*Sample Cost model: prior beliefs and risk attitude both improve the model fit.* The repeated trust game literature suggests that subjective prior beliefs (Chang et al., 2010) and betrayal aversion (Aimone & Houser, 2012) both play important roles in determining individual differences in trust. In our paradigm, an individual's prior belief about trustworthiness is a beta distribution with two free parameters ($\alpha_0$ and $\beta_0$). Betrayal attitude is operationalized as the variance of the outcome, multiplied by a free parameter $\lambda$, which is subtracted from the expected utility of trusting. The median betrayal attitude parameter was estimated at 0.16 (bootstrapped 95% CI [0.10, 0.23]), which shows that participants were overall betrayal-averse.

*Threshold model: asymmetric bounds but not collapsing bounds improved the model fit.* The literature on Drift Diffusion Models in perception literature suggests that the model fits better with collapsing bounds that reflect an urgency signal (Tajima et al., 2016), or asymmetric bounds (mulder et al., 2012) as positive sample outcomes might be weighted differently than negative outcomes. Using separate bounds for positive than for negative samples did indeed improve the model fit. Second, we tested the model when the bounds "collapse" to zero over time. This did not improve the model fit from the model with asymmetric bounds.

Table S1.1. Within-model comparison results

| | | 95% CI | |
|---|---|---|---|
| | Summed ΔBIC | Lower bound | Upper bound |
| Uncertainty basic vs. priors | 2013 | 925 | 3316 |
| Sample Cost basic vs. risk attitude | 1597 | 1006 | 2293 |
| Sample Cost risk attitude vs. risk attitude + priors | 791 | 398 | 1205 |
| Threshold basic vs. two bounds | 44987 | 38479 | 51847 |
| Threshold collapsing bound vs two bounds | 847 | 501 | 1198 |

95% CI = Bootstrapped 95% confidence interval of the summed difference between model fits. Smaller BIC values indicate better fit. Thus, positive values indicate a better fit for the second model. The models were fitted to the data at the individual level using a log likelihood optimization algorithm as implemented in the fmincon routine in MATLAB (©Mathworks). The optimization was iterated 100 times with varying initiations to avoid local minima. Because of the summation of the difference, large positive or negative numbers therefore reflect that one model wins consistently, i.e. for most subjects.

**Between- model comparison results**

We then compared the best fitting version of each model with the best fitting model of the other models. Table S2 shows the results which indicate that the Uncertainty model fitted best.

Table S1.2. Between model comparisons

| Pairwise comparison between models | Summed ΔBIC and 95% CI | Winning model | Correlation between age and ΔBIC |
|---|---|---|---|

| | | | |
|---|---|---|---|
| Uncertainty - Sample Cost | -1685 [-3086, -527] | **Uncertainty** | $r_s$ = -0.059, $P$ = 0.464 |
| Uncertainty - Threshold | -1741 [-3072, -636] | **Uncertainty** | $r_s$ = 0.101, $P$ = 0.221 |
| Sample Cost - Threshold | -56 [-627, 504] | **None** | $r_s$ = 0.031, $P$ = 0.711 |

Lower BIC values indicate a better fit, thus showing that the Uncertainty model fits significantly better than all other models. BIC scores were computed for each participant and each model. The BIC scores of a model pair (left column) was then subtracted from each other, thereby obtaining one difference score per participant for each model pair. The middle column shows the sum of the difference across participants and the 95% confidence interval of the BIC difference, computed using bootstrapping. For the winning model, we computed the correlation between age and each model pair's BIC difference. There was no significant correlation with age, suggesting that the best fitting model did not depend on age (right column). We report Spearman-rank correlations.

**Descriptive statistics**

The descriptive statistics were obtained through generalized linear mixed models in R (package LME4). For transparency we report the full regression in R code here:

```
SamplingModel <- glmer(Sample decision ~ |green – red| * poly(Age, degree = 2, raw = TRUE) + (1| subject), data = sampledata, family = binomial, control = glmerControl(optCtrl = list(maxfun = 1e+9), optimizer = c("bobyqa")))
```

This returned the following results:

Table S2.1. Results sampling decisions mixed model

| | Sample decisions | | | |
|---|---|---|---|---|
| *Predictors* | *Odds Ratios* | *95% CI* | *β* | *p* |

| | | | | |
|---|---|---|---|---|
| Intercept | 0.05 | 0.04 – 0.06 | -31.01 | **<0.001** |
| \|green – red\| | 2.29 | 2.22 – 2.37 | 52.29 | **<0.001** |
| Age linear | 1.05 | 0.93 – 1.18 | 0.73 | 0.465 |
| Age quadratic | 1.05 | 0.91 – 1.20 | 0.66 | 0.507 |
| \|green – red\| x Age linear | 1.19 | 1.17 – 1.21 | 16.79 | **<0.001** |
| \|green – red\| x Age quadratic | 0.96 | 0.94 – 0.99 | -3.05 | **0.002** |
| **Random Effects** | | | | |
| $\sigma^2$ | 3.29 | | | |
| $\tau_{00\ subject}$ | 0.59 | | | |
| ICC $_{subject}$ | 0.15 | | | |
| Observations | 162145 | | | |
| Marginal $R^2$ / Conditional $R^2$ | 0.137 / 0.268 | | | |

Similarly, the invest decisions were analysed as follows:

InvestModel <- glmer(Invest decision ~ reciprocation probability * poly(age, degree = 2, raw = TRUE) + (1| subject), data = investdata, family = binomial, control = glmerControl(optCtrl = list(maxfun = 1e+9), optimizer = c("bobyqa")))

This returned the following results:

Table S2.2. Results invest decisions mixed model

| | Invest decisions | | | |
|---|---|---|---|---|
| *Predictors* | *Odds Ratios* | *95% CI* | *β* | *p* |
| Intercept | 0.00 | 0.00 – 0.00 | -28.65 | **<0.001** |
| Reciprocation probability | 52426.19 | 28318.14 – 97058.12 | 34.58 | **<0.001** |
| Age linear | 0.35 | 0.27 – 0.45 | -8.02 | **<0.001** |
| Age quadratic | 1.65 | 1.24 – 2.18 | 3.49 | **<0.001** |

| | | | | |
|---|---|---|---|---|
| Reciprocation probability x age linear | 8.35 | 5.63 – 12.40 | 10.54 | **<0.001** |
| Reciprocation probability x age quadratic | 0.40 | 0.27 – 0.61 | -4.34 | **<0.001** |
| **Random Effects** | | | | |
| $\sigma^2$ | 3.29 | | | |
| $\tau_{00\ \mathrm{subject}}$ | 0.70 | | | |
| ICC $_{\mathrm{subject}}$ | 0.18 | | | |
| Observations | 9420 | | | |
| Marginal $R^2$ / Conditional $R^2$ | 0.753 / 0.797 | | | |

**Trust decisions**

Bonferroni-Holm corrected linear mixed models (alpha = 0.008) per reciprocation probability, showed that the age difference in trust decisions was only significant in the higher values of the generative reciprocation probability (i.e., when *r* was 0.8 and 1.0). Specifically, compared to older participants, younger participants trusted less often when the reciprocation probabilities were 0.8 ($\beta = 0.620$, $P < 0.001$) and 1.0 ($\beta = 0.922$, $P < 0.001$).

**Expected reward increased with age**

We examined if the expected values based on the trust decisions varied with age. We first normalized the outcomes on the endowment of 6 tokens. Thus, the expected value for not investing is 1. The expected value for investing is computed as the multiplier of 2 times the reciprocation probability. Since the reciprocation probabilities were 0.0, 0.2, 0.4, 0.6, 0.8, and 1.0, the possible average expected values ranged from $\frac{0 + 0.4 + 0.8 + 1 + 1 + 1}{6} = 0.67$ to $\frac{1+1+1+1.2+1.6+2}{6} = 1.3$. All subjects (with exception of one), had an average expected reward higher than 1 (see Figure S1). This confirms that all subjects would - on average - have gained money on top of their original endowment by trusting when trusting was beneficial. However, younger adolescents earned less on this task than older adolescents, consistent with their deviations in low (0.0) and high (0.8, 1.0) investment probabilities. This was shown by a significant correlation between age and the average expected reward in the task ($r_s = 0.387$, $P < 0.001$).

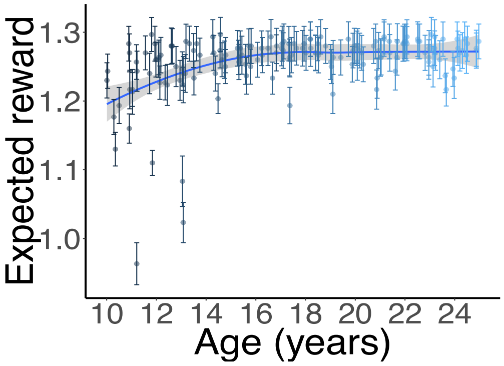


**Figure S1**. Expected reward per subject as function of age. The expected reward increases with age. Errorbars indicate the s.e.m. The line and shaded region indicates the mean and s.e.m across subjects. Color indicates age in months

Table S2.3. Post-hoc analyses of age effect on invest decisions per reciprocation probability

| | **Reciprocation probability = 0.0** | | | **Reciprocation probability = 0.2** | | | **Reciprocation probability = 0.4** | | | **Reciprocation probability = 0.6** | | | **Reciprocation probability = 0.6** | | | **Reciprocation probability = 1.0** | | |
|---|---|---|---|---|---|---|---|---|---|---|---|---|---|---|---|---|---|---|
| *Predictors* | *Odds Ratios* | *95% CI* | *p* | *Odds Ratios* | *95% CI* | *p* | *Odds Ratios* | *95% CI* | *p* | *Odds Ratios* | *95% CI* | *p* | *Odds Ratios* | *95% CI* | *p* | *Odds Ratios* | *95% CI* | *p* |
| Intercept | 0.00 | 0.00 – 0.01 | **<0.001** | 0.01 | 0.00 – 0.02 | **<0.001** | 0.17 | 0.14 – 0.22 | **<0.001** | 1.86 | 1.53 – 2.27 | **<0.001** | 20.95 | 14.76 – 29.75 | **<0.001** | 64.66 | 35.23 – 118.69 | **<0.001** |
| Age | 0.37 | 0.19 – 0.72 | **0.004** | 0.63 | 0.39 – 1.04 | 0.071 | 0.91 | 0.74 – 1.11 | 0.352 | 1.10 | 0.90 – 1.34 | 0.348 | 1.93 | 1.46 – 2.56 | **<0.001** | 2.88 | 1.88 – 4.40 | **<0.001** |
| **Random Effects** | | | | | | | | | | | | | | | | | | |
| $\sigma^2$ | 3.29 | | | 3.29 | | | 3.29 | | | 3.29 | | | 3.29 | | | 3.29 | | |
| $\tau_{00}$ | 5.03 $_{subject}$ | | | 4.70 $_{subject}$ | | | 0.86 $_{subject}$ | | | 1.06 $_{subject}$ | | | 1.01 $_{subject}$ | | | 1.83 $_{subject}$ | | |
| ICC | 0.60 $_{subject}$ | | | 0.59 $_{subject}$ | | | 0.21 $_{subject}$ | | | 0.24 $_{subject}$ | | | 0.24 $_{subject}$ | | | 0.36 $_{subject}$ | | |
| Observations | 1570 | | | 1570 | | | 1570 | | | 1570 | | | 1570 | | | 1570 | | |
| Marginal $R^2$ / Conditional $R^2$ | 0.108 / 0.647 | | | 0.025 / 0.599 | | | 0.002 / 0.209 | | | 0.002 / 0.246 | | | 0.092 / 0.305 | | | 0.179 / 0.472 | | |

### Age-related changes in Uncertainty model parameter estimates

Linear regressions were used to examine the relationship between age and the Uncertainty model parameter estimates using the lm() function in R. We first tested whether adding age as a quadratic term with the poly() function improved the model fits when compared to a linear term only using the anova() function in R (Phillips, 2017). For the prior uncertainty the quadratic term did improve the model fit ($F(1,154) = 6.835$, $p = 0.010$). For all other parameter estimates the quadratic effect of age did not improve the model fit (prior mean $F(1,154) = 3.861$, $p = 0.051$; uncertainty tolerance $F(1,154) = 1.940$, $p = 0.166$; decision noise $F(1,154) = 2.255$, $p = 0.119$). We applied a Bonferroni-holm correction for multiple testing, which includes the model improvement test for each of the four parameters, a linear age effect test for three of these parameters, and two test (linear and quadratic age effects) for one parameter, corresponding to a total of 9 tests.

### Model recovery

We performed model recovery and verified that the models were distinguishable. To this end, we simulated data with each model and subsequently fitted the simulated data to each model. If the model is recoverable, then the best fitting model should be the model that generated the data. We show that this was indeed the case as the model fit was always best for the model that generated the data (Table S2).

Table S3.1. Model recovery results

| | | 95% CI | |
|---|---|---|---|
| | Summed ΔBIC | Lower bound | Upper bound |
| **Data generated by the Uncertainty model** | | | |
| Uncertainty vs. Sample Cost | -849 | -1057 | -654 |
| Uncertainty vs. Threshold | -568 | -861 | -293 |
| **Data generated by the Sample Cost model** | | | |
| Sample Cost vs. Uncertainty | -795 | -313 | -541 |
| Sample Cost vs. Threshold | -537 | -908 | -249 |
| **Data generated by the Threshold model** | | | |
| Threshold vs. Sample Cost | -827 | -964 | -689 |
| Threshold vs. Uncertainty | -654 | -894 | -448 |

95% CI = Bootstrapped 95% confidence interval of the summed difference between model fits. Negative values indicate a better fit of the model that generated the data. The data were generated using the participants' parameter estimates and shows that all models were recoverable.

#### 3.2 Parameter recovery

To check if the parameters were recoverable, we simulated data with each model using varying parameter values as inputs. We randomly selected the estimates of 20 subjects to simulate data with an equal number of trials as in the actual task. We then fitted that simulated data to the model to check if the parameters values that generated the data were correctly estimated. We subtracted the estimated parameters of the simulation from the actual input parameters and calculated the bootstrapped 95% confidence interval and found no significant difference between the input parameters and the estimated parameter values. This returned: Uncertainty criterion median difference = -0.0001, 95% CI [-0.000, 0.000]; $\alpha_0$ median difference = -0.386, 95% CI [-0.590, 0.000]; $\beta_0$ median difference = -0.250, 95% CI [-0.580, 0.000]). This suggests that the parameters were indeed recoverable.
